# Biophysical models of protein evolution: Understanding the patterns of evolutionary sequence divergence

**DOI:** 10.1101/072223

**Authors:** Julian Echave, Claus O. Wilke

## Abstract

For decades, rates of protein evolution have been interpreted in terms of the vague concept of “functional importance”. Slowly evolving proteins or sites within proteins were assumed to be more functionally important and thus subject to stronger selection pressure. More recently, biophysical models of protein evolution, which combine evolutionary theory with protein biophysics, have completely revolutionized our view of the forces that shape sequence divergence. Slowly evolving proteins have been found to evolve slowly because of selection against toxic mis-folding and misinteractions, linking their rate of evolution primarily to their abundance. Similarly, most slowly evolving sites in proteins are not directly involved in function, but mutating them has large impacts on protein structure and stability. Here, we review the studies of the emergent field of biophysical protein evolution that have shaped our current understanding of sequence divergence patterns. We also propose future research directions to develop this nascent field.

## 1. INTRODUCTION

The success of theoretical physics in the 19th and 20th century has been driven primarily by the development of fundamental and elegant mathematical theories that accurately describe and predict physical processes from first principles. In biology, by contrast, development of fundamental theory has been lacking. Biology is much more a science of individual special cases and historical contingencies, and it has shown surprising resilience against physicists’ desire to describe it with theories that are both simple and general. However, the one area where biology may be most amenable to the development of fundamental theory is the field of protein evolution. Evolutionary biology is probably the most mathematical branch of biology, and sophisticated mathematical machinery exists to describe how populations accumulate mutations and evolve over time. Similarly, protein biophysics is governed by equilibrium and non-equilibrium thermodynamics, both well-developed areas of theoretical physics. In recent years, several groups have made efforts to combine evolutionary theory and protein biophysics, with the ultimate goal of developing a general, mechanistic theory of protein evolution.

Here we review both fundamental concepts and recent advances in the field of protein evolution. Since this field is extensive, no single review can cover it exhaustively. Recent reviews have covered empirical approaches to studying protein evolution, specifically reviewing the causes of rate variation among proteins (90) and among sites within proteins (21). Other reviews have emphasized how protein biophysics shapes fitness landscapes (14, 70, 74). To complement these works, we describe how theoretical population genetics and protein biophysics have been combined into predictive models of evolutionary rate variation among and within proteins. Our emphasis is on the development of theory, and we attempt to provide a comprehensive and self-contained description of the key concepts, starting from first principles.

### protein genotype

The amino-acid sequence of a given protein

### sequence space

Possible amino-acid sequences connected by single amino acid point mutations

### fitness

Expected number of offspring of an individual

### fitness landscape

Map from sequence space to fitness

## 2. PROTEIN EVOLUTION ON FITNESS LANDSCAPES

The evolution of a protein’s amino acid sequence (its genotype) can be represented as a trajectory in sequence space. Since evolutionary dynamics depends on fitness, protein evolution can be pictured as following paths on a fitness landscape (84). This section provides a basic introduction to the theory of evolution on fitness landscapes.

### 2.1. Population dynamics on fitness landscapes

Finite populations of genotypes evolve on the fitness landscape under genetic drift, selection, and mutation. The population’s evolution can be modeled by randomly drawing the genes of the offspring population from the pool of parent genotypes. Such stochastic sampling is called genetic drift. To model selection, individuals are drawn with weights proportional to their fitness. The drawn individuals may be mutated to introduce new genotypes.

A frequently used sampling model is the Moran process: For a population of haploid individuals of effective size *N*_e_, at each time-step one a random individual is removed and another individual, drawn randomly with a probability proportional to its fitness, is duplicated. Another popular model is the Wright-Fisher process of discrete non-overlapping generations: The offspring generation is obtained by *N*_e_ random draws with replacement from the parent generation, with probability proportional to fitness.

#### effective population size (*N*_e_)

Number of individuals that may contribute offspring to the next generation

In finite populations, stochastic fluctuations of allele frequencies result in genes becoming lost from the gene pool. If no new alleles are introduced, eventually all alleles but one will be lost and one allele will become fixed. If the time between introduction of new mutations is much larger than the timescale of fixation, most of the time all individuals of the population share the same allele (the population is monophyletic). Given a parent allele *i*, when a new mutant allele *j* is introduced it eventually is either lost or becomes fixed. For a haploid population evolving under the Moran process, the fixation probability is (44, 66):

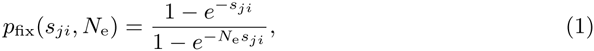

#### allele

Variant of a gene sequence generated through mutation

#### fixation

An allele is said to be fixed if it is present in all members of the population

#### fixation probability (*p*_fix_)

Probability of an allele becoming fixed

#### selection coefficient (*s_ji_*)

Fitness of allele *j* relative to allele *i*

where *s_ji_* is the selection coefficient (66):

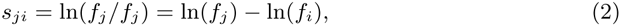

where *f_j_* and *f_i_* are the fitness values of alleles *j* and *i*, respectively. The selection coefficient measures the selective advantage of the mutant relative to the parent: *s_ji_* is zero for neutral mutations and positive (negative) for advantageous (deleterious) mutations. For the haploid Wright-Fisher process, *p*_fix_ can be obtained replacing *s_ji_* by 2*s_ji_* in equation 1. In certain conditions, fixation probabilities for diploid populations may be obtained replacing *N*_e_ by 2*N*_e_ (44).

#### origin-fixation model

Model of evolution that keeps track of only the most abundant genotype in the population; this genotype is assumed to change over time through the origination of individual mutations and their subsequent fixation in the population.

### 2.2. Origin-fixation models

For long time scales, evolution is often studied using origin-fixation models (46). Evolution is modeled using a Markov process that is completely specified by a rate matrix Q with off-diagonal (*j* ≠ *i*) elements:

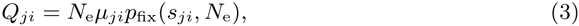

where *i* is the parent genotype, *j* the mutant, µ_*ji*_ is the *i* → *j* mutation rate per individual, and *p*_fix_ is the fixation probability of *j*. *Q*_*j*≠*i*_ represents the rate of *i → j* transitions; the diagonal elements satisfy *Q_ii_* = −Σ_*j≠i*_ *Q_ji_*.

In general, the time-dependent state of the system can be described by a (column) probability vector *π*(*t*) = (*π*_1_(*t*), π_2_(*t*), …)^*T*^, where π_*i*_(*t*) is the probability that at time *t* the fixed genotype is *i*. The evolutionary dynamics obeys:

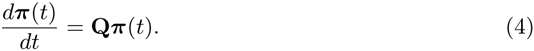

Eventually, *π*(*t*) approaches an equilibrium distribution of genotypes *π*\*. For origin-fixation models given by equation 3, with fixation probability given by equation 1 and a mutational process that satisfies 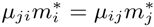, following (66) it can be found:

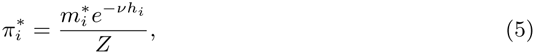

where 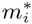 is the equilibrium probability of *i* under mutation only, *h_i_* = −ln(*f_i_*), *Z* = 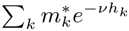, and *ν* = *N*_e_ − 1 for the haploid Moran process. (We have *ν* = 2(*N*_e_ − 1) for the haploid Wright-Fisher process and the diploid cases can be obtained by replacing *N*_e_by 2*N*_e_).

Note the mathematical similarity between equation 5 and the Boltzmann distribution of statistical physics, with 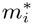 playing the role of the degeneracy of state *i*, *h_i_* = −ln(*f_i_*) the role of energy, and *ν* that of the inverse temperature *β*. This mathematical analogy is useful to import methods and concepts from statistical physics into molecular evolution, but it can also lead to confusion.

#### substitution rate

The rate at which mutations fix in the population per unit time

### 2.3. Substitution rate, entropy, and number of states

For further reference, we define here some quantities that reflect the degree of evolutionary constraint on protein sequences or on individual protein sites. Consider a distribution of genotypes *π*. One measure of constraint is the substitution rate *K*, which is the mean number of amino-acid substitutions per unit time:

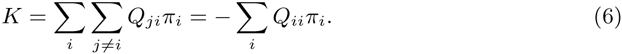

Another measure of constraint is sequence entropy, which quantifies how dispersed the distribution of genotypes is:

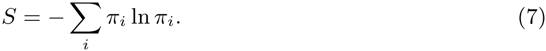

Often, a more convenient measure is

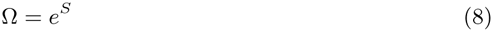

which is a number between 1 and the total number of states; it represents the effective number of different states allowed.

## 3. MOLECULAR, BIOPHYSICCS-BASED FITNESS LANDSCAPES

The key challenge in developing a biophysical description of protein evolution is to determine how fitness f depends on the amino-acid sequence. A mutation will impact the fitness of the organism in relationship to how the mutation (i) perturbs the activity of the protein and (ii) creates any undesired side-effects on the organism. For example, to be functional, the protein may have to be able to fold rapidly into a given specific native structure, bind some substrate, and so on. Similarly, to not create side effects, a protein must not interact with off-target binding partners or create toxic metabolic products. These traits (structure, folding, stability, binding affinity, enzymatic activity, etc.) constitute the molecular phenotype of the protein, those protein properties that affect organism fitness.

### 3.1. Thermodynamics of protein folding

According to statistical thermodynamics, at constant pressure and temperature, the probability *P_i_* to observe a protein in a given conformation *i* is given by the Boltzmann distribution,

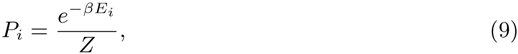

where *E_i_* is the potential energy of conformation *i* and β = (*k_B_T*)^−1^ (with *k_B_* being the Boltzmann constant and *T* the physical temperature of the system). The quantity *Z* = Σ_*j*_ *e*^−β*E_i_*^ is called the conformational partition function. The energy *E_i_* itself is calculated from some potential energy function, e.g., a pairwise potential summed over all pairs of atoms in the conformation *i*.

In practice, we are usually interested in the ground-state of the protein. The ground state will typically correspond to a set of conformations with low energy, and we also refer to it as the folded state. The probability to observe the folded state, *P*_folded_, relative to the probability to observe any other state, *P* unfolded, also follows from a Boltzmann distribution:

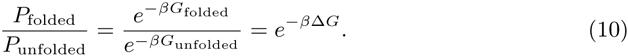

The quantity Δ*G* = *G*_folded_ − *G*_unfolded_ is the free energy of folding, and it measures the stability of the folded protein. The more negative Δ*G*, the more stable the protein. The stability Δ*G* of a protein will typically change if the protein is mutated. In many modeling scenarios, we are interested in the magnitude of this change, commonly denoted by ΔΔ*G*.

Equation 9 (describing the probability to observe a specific protein in a particular conformation) looks similar to equation 5 (describing the probability to observe the evolutionary trajectory visit an individual genotype). However, these two equations describe two entirely distinct processes and one does not follow from the other. To relate equation 9 to biological evolution, we need to introduce an explicit mapping to fitness, see Section 3.2.

### 3.2. Fitness landscapes based on protein fold stability

Most biophysical models of protein evolution map stability Δ*G* to fitness. Several examples are illustrated in Figure 1. For example, if fitness is assumed to be proportional to the probability of folding, then we obtain the soft-threshold model shown in **Figure 1*a*** (82):

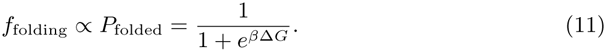

**Figure 1.**
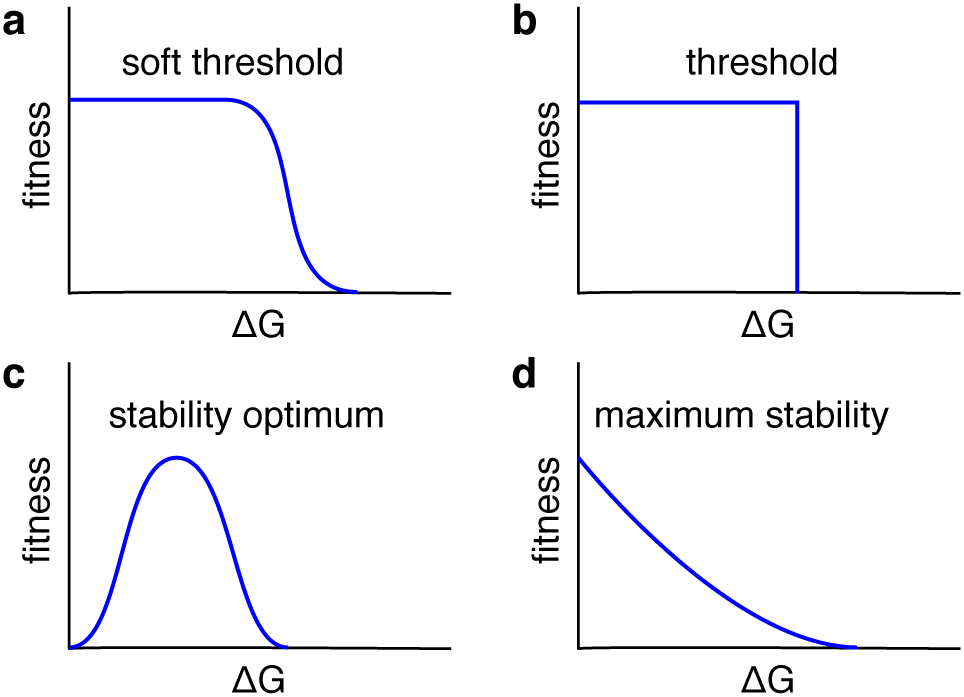
Common stability-based fitness models. (*a*) The soft-threshold model assumes that sufficiently stable proteins (low Δ*G*) all have the same high fitness, whereas unstable proteins (high Δ*G*) have fitness zero. Inbetween these two extremes, there is a smooth, sigmoidal transition region. (*b*) The threshold model can be considered as a limiting case of the soft-threshold model. It assumes that all proteins with sufficient stability (Δ*G* < some threshold) have the same fitness, while all other proteins are completely inviable. (*c*) The stability-optimum model assumes that there is an ideal stability for maximum fitness, and both more and less stable proteins will be less fit. (*d*) The maximum-stability model assumes that fitness is the higher the more stable the protein.

A simplified version of the soft-threshold model uses a hard stability threshold (**Figure 1*b***) (6, 7, 20). This model can be considered the limit of the soft-threshold model for very large β (low physical temperature). Alternatively, if protein stability and protein activity trade off, then we expect fitness to be maximal at some intermediate optimum stability (**Figure 1*c***) (14, 71). Finally, if fitness declines with increasing amounts of misfolded protein in the cell, due to protein toxicity, then we obtain the maximum-stability model of **Figure 1*d*** (18, 69, 86):

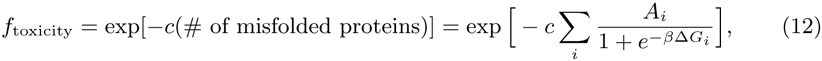

where *A_i_* is the abundance of protein *i*, *c* is a measure of toxicity per misfolded protein, and the sum runs over all genes *i* in the organism’s genome.

There are several modeling approaches that are commonly used to calculate the stability Δ*G* for a given protein sequence. First, we can evaluate equation 9 directly for models in which we can exhaustively enumerate all possible energy states *E_j_* (74). Second, we can estimate *Z* from random energy models (22, 73) or ignore it entirely (42). These approaches assume that individual mutations have a negligible effect on *Z*. Alternatively, a more realistic approach is to estimate *Z* from a limited number of decoy structures (82). Some authors also use detailed, all-atom off-lattice models (2, 15, 20, 37, 71), but to date this approach has seen limited use in evolutionary modeling, due to its high computational cost. Finally, some models operate entirely in Δ*G* space, keeping track of how Δ*G* changes due to the stability effects ΔΔ*G* of successive mutations (69).

### 3.3. Fitness landscapes based on protein function

Stability-based models such as the soft-threshold or stability-optimum models are implicitly modeling protein function, by assuming that either all folded proteins or only the proteins with the right optimum stability are functional. However, how exactly a protein’s function contributes to organism fitness is highly specific, and hence in principle we would have to derive function-based fitness landscapes from the biochemical properties and interactions of each individual protein. However, there are at least two broad classes of models that can be used to describe a generic, functional protein.

First, many authors have modeled function through protein–protein or protein–ligand binding (9, 31, 37, 49, 59, 83, 85). For example, fitness can be assumed to be proportional to the binding energy of the protein–ligand complex or the probability of finding the protein in the bound state. In all cases, the probability of binding is measured through a change in free energy Δ*G*_binding_, which is calculated according to the same principles of thermodynamics outlined in Section 3.1. For further details, see e.g. Refs. (49, 83).

Second, for enzymes, we can consider the flux through metabolic pathways (67). The fitness contribution due to flux through a linear pathway can be written as (67)

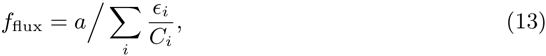

where *a* is the abundance of the input metabolite, ϵ_*i*_ is the enzymatic activity of protein *i* in the pathway, and *C_i_* is the number of functional copies of protein *i*. *C_i_* in turn can be written as the protein expression level *A_i_* times the probability that protein *i* is folded (equation 11), yielding the expression

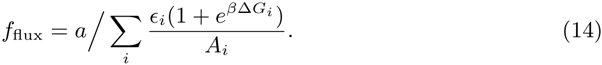

This equation describes how *f* _flux_ depends on protein stability. It does not, however, explicitly model how the proteins’ enzymatic activities ϵ_*i*_ depend on mutations.

Models of ϵ_*i*_ for actual proteins are still lacking. However, enzyme-like constraints have been considered explicitly in studies with simplified protein-like polymers. A model by Sasai and coworkers assumed that fitness increases monotonously as the conformation of a few active site residues converges to a predefined active conformation (62, 63, 88). Similarly, a recent study used a threshold-like fitness function that constrains the structure of a ligand and a few sites that bind it (51).

## 4. MODELING AMONG-PROTEIN RATE VARIATION

We will now discuss models that combine molecular fitness landscapes with models of sequence evolution to calculate the rates at which proteins accumulate substitutions. These models attempt to explain why some proteins evolve faster or slower than others. They typically consider only the average rate of evolution across the entire protein and neglect variation among sites. Variation among sites may be implicitly present in the model, but this variation is not the focus of the analysis and it is averaged out in the final modeling predictions.

### 4.1. More robust protein structures evolve more rapidly

First, we can ask how rapidly a protein evolves if it is only required to fold stably into its native conformation (the threshold or soft-threshold models of **Figure 1**). Under this model, we would expect that proteins whose structures are, on average, more tolerant to substitution (i.e., more mutationally robust) should evolve more rapidly (5). However, in practice, it is not entirely clear how to measure the robustness of a protein structure. A random-energy model predicts that a structure’s robustness is determined by its contact matrix (22). Based on this prediction, the average number of residue–residue contacts, the principal eigenvalue of the contact matrix, and the size of the protein core (as measured e.g. by the fraction of buried residues) have all been proposed as simple measures of mutational robustness (5). And indeed, there is evidence that proteins with higher contact density or with a larger core evolve more rapidly (5, 23, 26, 27, 64). Alternatively, it should be possible to calculate a protein’s robustness to mutations from its ΔΔ*G* distribution (8, 80). However, this approach has been less successful at explaining observed rate variation among proteins (23).

Importantly, the mutational robustness of a protein changes as the protein mutates, and the evolutionary rate is determined by the average robustness of all sequences fixed during the evolutionary trajectory. This average robustness will depend on intrinsic properties of the structure (some structures are, on average, inherently more robust to mutations than others) as well as extrinsic factors such as mutation rate *µ* and population size *N*_e_. If the product of mutation rate and population size is large, *µN*_e_ ≫ 1, then populations concentrate in the most connected (i.e., most mutationally robust) regions of the protein neutral network (10, 78). Therefore, as we increase *µ* or *N*_e_ or both, the mean substitution rate increases as well (7, 79).

#### protein neutral network

The network of protein sequences that fold stably into the native conformation of the protein

### 4.2. Proteins evolve to be robust against misfolding

One of the best predictors of among-protein evolutionary rates, in many different systems, is the cellular abundance of proteins. The more highly expressed a protein is, the slower it evolves. Because of both the strength and universality of this effect, it requires an explanation that applies in any organism, is consistent with protein biophysics, and is capable of reproducing the key patterns of sequence variation observed in natural proteins. The first explanation that met these criteria was the idea that organisms experience strong selection pressure against protein misfolding, specifically against misfolding due to errors introduced during translation (17).

Fitness due to misfolding is generally assumed to decline exponentially in the number of misfolded proteins, as in equation 12 (18, 81). This number can be calculated, for example, via a lattice-protein simulation that simulates error-prone translation (18). The lattice-protein model produces a power-law relationship between evolutionary rate and abundance, as observed in natural sequences, and it also reproduces observed codon-level selection pressures, such as an increase in the fraction of preferred codons for more highly expressed genes. While this model only considers misfolding induced by translation errors, proteins may also misfold spontaneously, without a proximate cause (19). A variant of the lattice-protein simulation allowing for spontaneous misfolding (86) has demonstrated that indeed the source of protein misfolding, spontaneous or induced by translation errors, is not critical to observe the key evolutionary adaptations that protect against misfolding.

However, the lattice-protein simulations used to date considered only a single protein fold; variation among folds was not considered (18, 86). Alternatively, Lobkovsky et al. estimated protein misfolding in a coarse-grained off-lattice model (42) and calculated the resulting rate distribution across folds. They found that their calculated rate distribution matched the observed distribution of rates in both bacteria and mammals. The major limitation of (42), however, was that variation in protein abundance was not considered.

As an alternative to using an explicit protein folding model, either on or off lattice, one can also describe the protein misfolding probability in terms of the ΔΔ*G* distribution (68, 69). This approach has the benefit of yielding analytic insight into the relationship between substitution rate, protein abundance, and population size. For the simplest such model, which assumes that the ΔΔ*G* distribution is constant and independent of stability Δ*G* (2, 39, 76), the mean substitution rate is independent of protein abundance (69). To recover a rate–abundance anticorrelation, mutations need to become increasingly destabilizing as protein stability increases (68, 69). This relationship between protein stability and the mutational effect of mutations is realized in all explicit folding models.

### 4.3. Proteins evolve to avoid misinteraction

Protein misfolding is not the only mechanism that creates a fitness cost proportional to protein abundance. Even if a protein is folded correctly, it may interact with off-target partners, and such interactions are likely deleterious. Since the probability of off-target interactions increases in proportion to the abundance of a given protein, it is reasonable to assume that the fitness cost due to such protein misinteractions similarly increases with abundance.

The importance of protein misinteractions was first pointed out by Zhang et al. (89), who estimated the amount of protein misinteractions in a cell and proposed an upper limit to gene expression and proteome size. Subsequently, Yang et al. (85) used a lattice-protein simulation, similar to the simulation models employed to study misfolding (18, 86), but now using a fitness function based on target and off-target interactions. In their model, fitness increases linearly with the proportion of correct interactions and decreases exponentially in proportion to the amount of non-specific interactions. This model produces a rate–expression-level anti-correlation, just like the models based on misfolding do. More importantly, it provides an explanation for the observation that there is a larger decrease in rate at solvent-exposed residues than at buried residues with increasing expression level (27, 64, 85).

Importantly, to date no model has combined both misfolding and misinteractions. Experiments support both effects, a fitness cost from protein misfolding (28) and widespread off-target interactions among proteins (13). Thus, the challenge for future work will be to determine the relative importance of these and other mechanisms.

### 4.4. Proteins evolve to carry out their function

While models based on protein stability, protein misfolding, and/or protein misinteraction reproduce major patterns of evolutionary-rate variation in natural proteins, they neglect a key component that must have some effect on rate variation: protein function (90). For example, a negative correlation between rate and abundance can also be explained by the expression cost hypothesis, which argues that the fitness reduction due to a deleterious mutation affecting protein function should scale with protein abundance (11, 30) [but see discussion in (90)].

In general, even though there are well-developed models of how fitness may depend on protein function (Section 3.3), few authors have used these models to calculate among-protein rate variation. One exception is the misinteraction model by Yang et al. (85), which also contains a fitness term for correct, functional interactions. However, both this fitness term and the one for misinteractions are ad hoc, not derived from first principles.

An alternative, promising direction was recently taken by Serohijos and Shakhnovich (67), who model the metabolic flux through a pathway and write fitness as the difference between the fitness components *f*_flux_, due to flux (equation 14), and *f*_toxicity_, due to toxicity caused by protein misfolding (equation 12). However, how these different fitness components interact in their effect on evolutionary rate has not yet been studied in detail.

## 5. MODELING AMONG-SITE RATE VARIATION

Not only do mean evolutionary rates vary among proteins, but rates also vary among sites within proteins. More generally, not only rates but the whole pattern of sequence divergence is site-dependent. In this section, we describe modeling efforts aimed at explaining among-site variation of sequence divergence patterns.

### 5.1. Evolution at the site level

Sequence divergence patterns at the site level can be obtained from simulations. For instance, suppose that we set up a model combining some evolutionary model of Section 2 with some molecular fitness landscape of Section 3 and we simulate divergent evolution by running several independent trajectories that start at the same ancestral protein sequence. Then, for any given site *l*, we may estimate its amino acid probability distribution *π*^*l*^ as the fraction of trajectories that have each amino acid at the focus site. Given *π*^*l*^, we may calculate site-specific sequence entropies *S^l^* and numbers of observed amino-acids *Ω^l^* (using equations 7 and 8, respectively). Counting amino acid substitutions along the trajectories, we can calculate site-specific substitution rates *K^l^* and even site-specific 20*×* 20 matrices of rates of amino acid substitutions *Q^l^*.

### 5.2. Site-specific and pooled analyses

A simple but often overlooked issue is that sequence patterns of individual sites are generally different from those of sites pooled together into groups. For instance, the variability of individual sites increases monotonically with relative solvent accessibility (RSA) (75), as shown in **Figure 2*a***. In contrast, the variability of groups of sites of similar RSA is much larger and has a peak at some intermediate RSA value, as shown in **Figure 2*b***. The difference between per-site analysis and pooled analysis is explained schematically in **Figure 3**.

**Figure 2.**
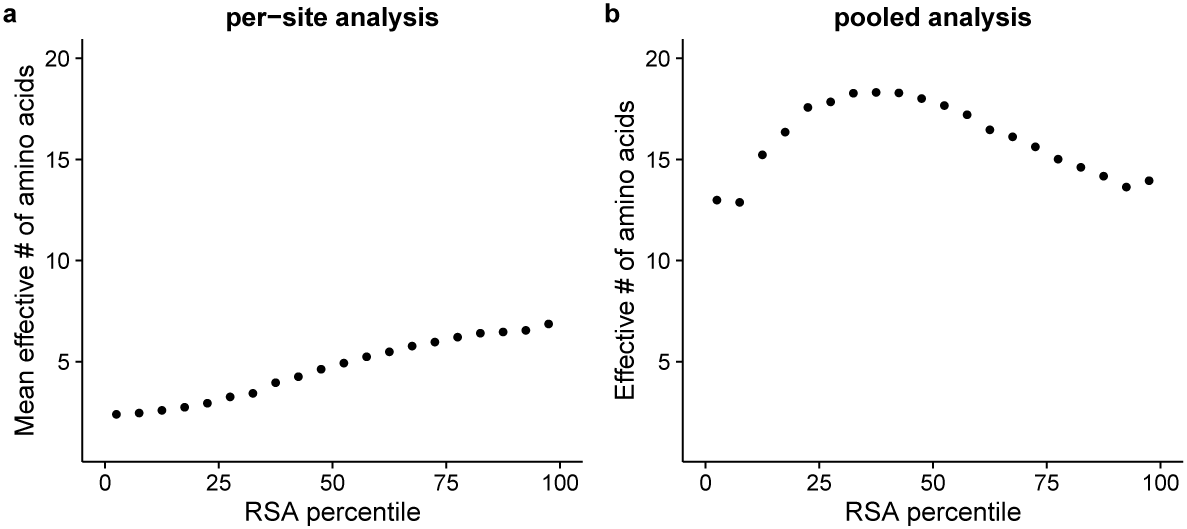
The relationship between the effective number of amino acids Ω and the relative solvent accessibility (RSA) of sites depends on whether amino-acid distributions are pooled across similar sites. (*a*) Without pooling, an Ω value is calculated at each site, and then Ω is averaged over all sites within the same RSA percentile. This procedure yields relatively low Ω values, between 2 and 10, and a nearly linear increase of Ω with increasing RSA. (*b*) Alternatively, amino-acid distributions are pooled among all sites within the same RSA percentile, and then a single Ω value is calculated for each pooled distribution. This procedure yields much higher Ω values (*>* 10), and it shows a maximum at intermediate RSA values. Protein sequences and structural data for this analysis were taken from (34). We selected 266 distinct enzymes for which each available alignment consisted of at least 400 sequences.

**Figure 3.**
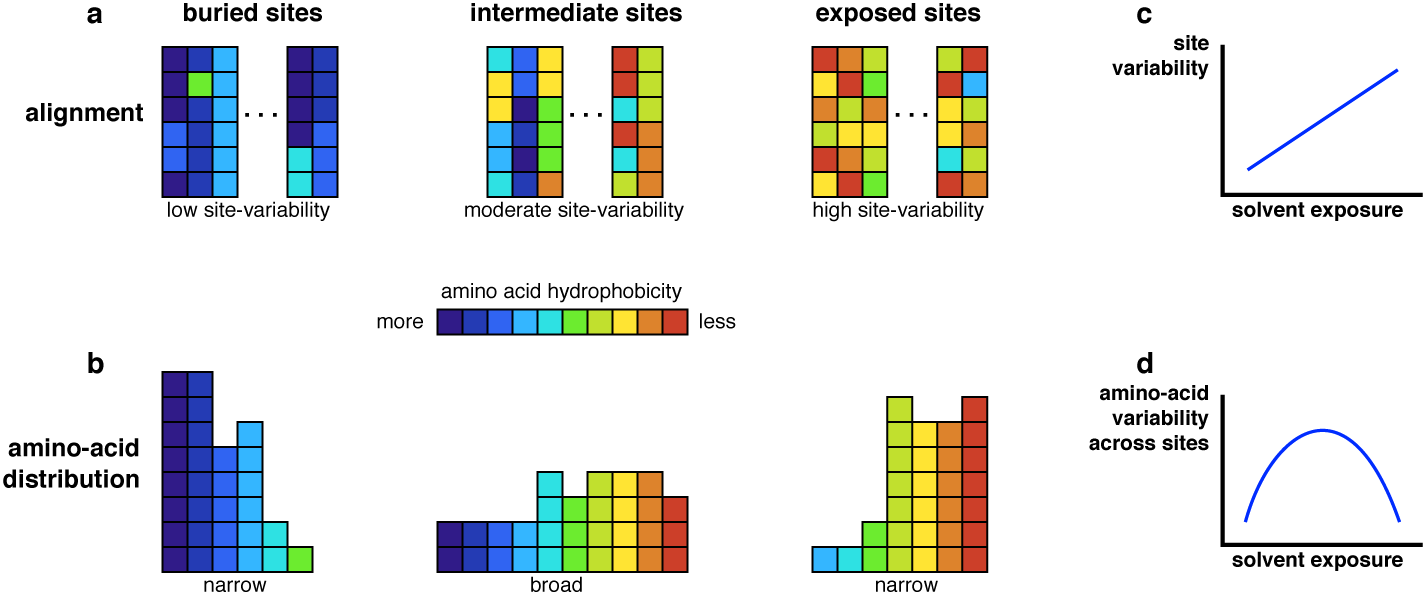
Conceptual explanation for the results shown in **Figure 2**. (*a*) At a per-site level, buried sites are most conserved, intermediate sites are moderately conserved, and exposed sites are the most variable. (*b*) However, for both buried and exposed sites, the types of amino acids that are seen are limited to mostly hydrophobic or mostly polar residues, respectively, whereas for intermediate sites both hydrophobic and polar residues are seen across many sites (58). As a consequence, even though the site variability increases approximately linearly with solvent exposure (*c*), the amino-acid variability across sites has a maximum at intermediate solvent exposure (*d*).

Models that are assessed using pooled analysis may fail to reproduce site-level patterns. For instance, a mean-field (MF) model by Bastolla and coworkers predicts that sites with intermediate solvent accessibility are the most variable, with more exposed sites having conservation levels similar to buried sites (3, 4, 57). This prediction is satisfied for pooled sites (**Figure 2*b***) but is violated for individual sites (**Figure 2*a***) (58).

### 5.3. Selection for rapid folding

It is reasonable to assume that since proteins need to fold rapidly, sites critical for folding kinetics should be particularly conserved. Early studies aimed at finding evidence of special conservation of protein sites involved in the folding nucleus (a few sites that are key for rapid folding) (48, 72). Starting at random sequences, sequence trajectories were run selecting for maximization of kinetic stability (the energy gap between the target conformation and the lowest-energy misfolded conformation). By comparing site-specific entropies of simulated and natural sequences, it was found that while folding nucleus sites were more conserved than average, many non-nucleus sites were similarly conserved due to stability constraints alone. Special conservation of the folding nucleus was later disproved by other authors (40, 77). This finding is consistent with kinetic stability correlating strongly with thermodynamic stability (50).

### 5.4. Selection on native state stability

Since most proteins need to fold stably into their native conformation to be functional and/or to avoid toxic effects of misfolded and unfolded conformations, it is reasonable to ask whether stability-based models explain among-site evolutionary variation.

Some authors have studied site-specific patterns modeling evolution using sequence design algorithms (15, 35, 52). Sequence design can be seen as a model of evolution with selection for stability (e.g. **Figure 1*d***), because sequences are designed to maximize the stability of a target structure. In general, the agreement between designed and natural sequences is moderate (35, 52). Specifically, predicted site-specific entropies increase too slowly with increasing relative solvent accessibility (RSA) and designed proteins have too few hydrophobic residues and too many hydrogen bonds in the core (35). The mismatch between designed and natural sequences may have several causes, such as errors in the energy function or the fitness function. More importantly, a major caveat of sequence design is that it simulates convergent evolution starting from random sequences, rather than divergent evolution starting from a common ancestral sequence. While the convergent and divergent processes should lead to the same equilibrium distributions at infinite time, natural sequences have diverged for a finite amount of time and finite-time sequence patterns may look different from the infinite-time steady state picture.

Divergence from a common ancestor with a fitness function with a stability optimum (**Figure 1*c***) is the basis of the Structurally Constrained Protein Evolution (SCPE) model (53, 55). This model simulates divergent evolution from a protein of known structure and sequence, which is assumed to be at the fitness peak. Mutations are introduced according to a DNA level process and selected against departure from the optimum. Site-specific amino-acid distributions, entropies, rates, and substitution matrices are calculated as explained in Section 5.1 (24). For a highly regular structure for which sites can be grouped into classes, the model predicted quantitatively class-specific amino acid distributions and entropies (53, 54). More generally, predicted site-specific substitution matrices fit homologous sequence families better than do site-independent models (55). Interestingly, a recent study showed that for many ligand-binding proteins with more than one stable conformation, the SCPE site-specific matrices obtained from the ligand-binding conformation fit sequence data better than matrices obtained from the inactive conformations (36).

In contrast with the simulation approach of the previous paragraphs, a recent study derived substitution-rates theoretically for a stability-threshold model (**Figure 1*b***) (20). This work built on a model by Bloom and Glassman (6) to derive an analytic relationship between mutational ΔΔ*G* values and site-specific substitution rates. ΔΔ*G* values for all possible single-point mutations were calculated for a large set of monomeric enzymes using FoldX (32, 65), and predicted rates were found to correlate well with empirical rates (20). This result shows that native state stability constraints can explain part of the observed rate variation among protein sites.

### 5.5. Selection on active structure stability

Since proteins fluctuate around the minimum energy conformation, the native state consists of an ensemble of conformations. The stability-based models of Section 5.4 are based on the total probability of the native state ensemble (e.g. equation 11). By contrast, a recently developed stress model assumes that fitness depends on the probability of specific active conformation (33, 43). Mutations shift the conformational ensemble, modifying the probability of reaching the active conformation. Basic statistical physics was used to derive an analytic relationship between site-specific rates and the impact of mutations on the stability of the active conformation, 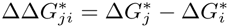.

In the published studies, ΔΔ*G*\* was calculated using perturbed elastic network models (33, 43). Amino acids were represented by nodes connected by harmonic springs and mutations were modeled as random perturbations of the spring-lengths of the mutated site. For a large set of monomeric enzymes, the stress model explained the observed linear decrease of site-specific substitution rates with increasing local packing density and their non-linear increase with site flexibility: The average destabilization of the active conformation is proportional to local packing density, and therefore tightly packed sites evolve more slowly than loosely packed ones (26, 33, 87). Moreover, the model predicted side-chain packing density to be the main structural determinant of a site’s rate of evolution (43).

### 5.6. Selection on protein function

The models described so far in Section 5 do not explicitly consider specific functional constraints. Therefore, it is often the case that some sites are observed to be much more conserved than predicted by such models. A few biophysical modeling studies found that including explicit functional constraints may affect site-specific sequence divergence.

One form of functional constraints are constraints induced by protein–ligand interactions. Only a few modeling studies have considered the effect of ligand-binding constraints on site-specific sequence divergence (31, 41, 59). Selection for binding affects the site-specific patterns of the binding site. Importantly, selection against binding non-target ligands needs to be taken into account to reproduce (qualitatively) the conservation patterns observed in natural sequences (31).

Protein–protein interactions may also affect divergence patterns. For multimeric proteins, SCPE model simulations that include monomer–monomer interactions improve the fit of site-specific substitution matrices for the sites directly involved in the interactions (25). However, the overall conservation of binding surfaces is low. A more recent study that included protein–protein interactions in simulations of sequence divergence found that binding constraints have surprisingly little influence on the rate of divergence of the sites involved in the binding surface (37). The binding surface diverges nearly as fast when selection for continued binding is imposed as when this selection pressure is removed. This finding is consistent with divergence patterns observed at binding interfaces in enzymes (34).

In contrast with the low conservation observed in protein-protein binding surfaces, active sites in enzymes are very conserved and impose long-range constraints on the evolution of most protein sites (34). This conservation is probably caused by the requirement of residues involved in catalysis to adopt very precise conformations in the substrate binding pocket. We lack realistic models of functional selection for enzyme evolution. However, the effect of constraining just a few sites on protein evolution has been studied using simple protein-like polymers. Sasai and coworkers simulated the evolution of protein-like polymers with a fitness function that constrains only the structure of a few active residues (62, 63, 88). Remarkably, they found that such functional constraints alone were enough for sequences to evolve into stable, ordered, protein-like molecules (62, 88) and that the rate of evolution of a site increases with its flexibility (63). Recently, Nelson and Grishin used a similar protein-like model to study the effect of enzyme-like functional constraints on sequence divergence more thoroughly (51). Starting from ordered structures, they simulated divergent evolution by introducing mutations, insertions, and deletions, constraining only the structure of a small binding pocket whose sites were not mutated. These simulations reproduce many recent findings: (i) site-specific rates increase linearly with solvent accessibility (26), (ii) rates decrease linearly with local packing density (33, 87), (iii) packing density is a better predictor of rates than solvent accessibility (43, 87), (iv) rates increase non-linearly with flexibility (33). Importantly, the model also predicts (v) a linear increase of site-specific rates with distance to the active site, in agreement with recent findings in a large dataset of over 500 enzymes (34).

### 5.7. Epistasis and time-dependence of site-specific patterns

So far, we have considered site-specific divergence patterns averaged over time. However, site-specific patterns may change along an evolutionary trajectory, because the selection coefficient of a given mutation may depend on the specific sequence background in which it occurs. This phenomenon, called epistasis, has been studied extensively in empirical and experimental research, but only recently has it been investigated using biophysical models of protein evolution (2, 56, 71).

Epistatic effects have been clearly established in an elegant study by Shah et al. (71). They simulated sequence evolution on two fitness landscapes, a (gaussian) stability optimum and a (semi-gaussian) soft threshold (see **Figure 1**). For both cases, they found that fixed mutations are nearly neutral and contingent on previous substitutions: The same mutation would be too deleterious to fix if previous permissive substitutions had not occurred. In addition, substitutions become entrenched over time: A mutation that was nearly neutral when fixed becomes increasingly deleterious to revert as subsequent substitutions accumulate.

Entrenchment had been previously demonstrated in a stability-based simulation study by Pollock et al. (56). They found that as a result of entrenchment, site-specific amino acid distributions are time-dependent. However, Bloom and coworkers have called into question the prevalence and magnitude of time-dependent epistatic shifts in site-specific distributions (2). Using simulations analogous to those of Pollock et al., they found that while site-specific amino-acid propensities do change, they diverge very slowly, so that they are mostly conserved among homologous proteins over realistic time scales.

The degree of variation of site-specific amino acid distributions with time depends on whether deleterious mutations are compensated by reversion or entrenchment. While entrenchment makes reversion increasingly unlikely (56, 71), reversions may be the most common way of compensating deleterious substitutions (2, 16). A step forward towards understanding the reversion–entrenchment trade-off is the recent finding that for a site involved in epistatic interactions, the probability of reverting a deleterious substitution is large just after fixation and then decreases rapidly with time (45).

That epistasis does play a role in protein evolution is widely accepted. However, the prevalence and magnitude of epistatic time-dependent propensity shifts and their implications for phylogenetic modeling of homologs with similar structures and functions is still under debate (16, 29).

## 6. CONCLUSIONS AND OUTLOOK

Theoretical population genetics studies evolution on fitness landscapes. Protein biophysics aims to understand protein function from the fundamental laws of physics. Insofar as fitness depends on protein function, protein biophysics should be able to derive the fitness landscape from first principles. In recent years, researchers have combined evolutionary models with fitness functions based on biophysical properties to derive biophysical models of protein evolution, giving rise to the emergent field of biophysical protein evolution.

Arguably, the main goal of a theory of protein evolution is to explain and predict patterns of amino acid sequence divergence. Biophysical evolution studies have already provided major advances in our understanding of why evolutionary rates vary among proteins, among sites within proteins, and over time. Regarding rate variation among proteins, the observed rate–expression anticorrelation may be due to several different biophysical mechanisms, including mistranslation-induced misfolding (18), spontaneous misfolding (69, 86), and misinteraction (85). Regarding variation among sites, the increase of rates with solvent accessibility and flexibility and its decrease with local packing density may be explained in terms of folding stability and/or stability of active conformations (21). Finally, variation of rates over time follows from epistatic interactions inherent in selection on stability (2, 56, 71).

Ideally, the fitness landscape should be derived from the biophysical principles that govern protein behavior. However, the fitness functions employed so far remain still mostly ad hoc. For example, a fitness function that posits an optimum stability corresponding to maximal fitness represents a biophysical model, but it does not provide any insight into whether or why fitness may be maximal at that stability value. By contrast, a soft threshold derived from *f* ˜ *P* folded can be considered more principled, since it represents the assumption that activity is proportional to the abundance of proteins in the native state. Therefore, one broad research direction is to develop more principled biophysical models of fitness.

We see several major avenues for future work. First, the existing models need to be studied more exhaustively, across a larger range of different scenarios. For example, most of the models that are critical to understanding variation of rates among proteins, such as models of mistranslation, misfolding, and misinteraction, have not been used to study site-specific patterns, even though these processes will likely likely affect site-specific patterns as well. Second, we need to develop better models of functional constraints. For instance, it is not yet clearly understood why enzyme active sites impose long-range evolutionary constraints on most protein sites. Third, how epistasis among protein sites affects the substitution process at individual sites remains poorly understood and deserves further study. Finally, biophysical models of protein evolution can be integrated directly into phylogenetic methods of sequence analysis, alleviating the need of having to first calculate summary statistics of the evolutionary process (such as rates or entropies) and then correlating these statistics with model predictions. While some progress has been made in this direction (1, 12, 38, 47, 60, 61, 64), most of the models proposed to date have been computationally cumbersome and ultimately not yet useful.

#### SUMMARY POINTS

1. Biophysical models of evolution combine population genetics models of evolution on fitness landscapes with biophysical models of the fitness landscape.
2. Most current biophysical models posit ad hoc fitness functions that depend on biophysical traits that are assumed to affect protein function or toxicity.
3. In spite of their mostly ad hoc nature, biophysical modeling studies have significantly advanced our understanding of why the rate of evolution varies among proteins, among sites within proteins, and over time.

#### FUTURE ISSUES

1. Derive less ad hoc fitness functions from first principles of molecular biophysics.
2. Develop fitness models that consider the joint effects of distinct biophysical mechanisms.
3. Develop better models of functional constraints, especially for enzymes.
4. Clarify how epistasis shapes evolutionary divergence at individual protein sites.
5. Integrate directly biophysical protein-evolution models into phylogenetic methods of sequence analysis.

## DISCLOSURE STATEMENT

The authors are not aware of any affiliations, memberships, funding, or financial holdings that might be perceived as affecting the objectivity of this review.

## ACKNOWLEDGMENTS

This work was supported in part by NIH grants R01 GM088344 and R01 AI120560, NSF Cooperative agreement DBI-0939454 (BEACON Center), and ARO grant W911NF-12-1-0390.

